# Guided Co-clustering Transfer Across Unpaired and Paired Single-cell Multi-omics Data

**DOI:** 10.1101/2025.05.16.654635

**Authors:** Hongyao Li, Yunrui Liu, Pengcheng Zeng

## Abstract

Single-cell multi-omics technologies enable the simultaneous profiling of gene expression and chromatin accessibility, providing complementary insights into cellular identity and gene regulatory mechanisms. However, integrating paired scRNA-seq and scATAC-seq data remains challenging due to inherent sparsity, technical noise, and the limited availability of high-quality paired measurements. In contrast, large-scale unpaired scRNA-seq datasets often exhibit robust and biologically meaningful cell cluster structures. We introduce **Guided Co-clustering Transfer (GuidedCoC)**, a novel unsupervised framework that transfers structural knowledge from unpaired scRNA-seq source data to improve both cell clustering and feature alignment in paired scRNA-seq/scATAC-seq target data. GuidedCoC jointly co-clusters cells and features across modalities and domains via a unified information-theoretic objective, aligning gene expression modules with regulatory elements while implicitly performing cross-modal dimensionality reduction to reduce noise. Additionally, it automatically aligns cell types across unpaired and paired datasets without requiring explicit annotations. Extensive experiments on multiple benchmark datasets demonstrate that GuidedCoC achieves superior clustering accuracy and biological interpretability compared to existing methods. These results highlight the promise of structure-guided, unsupervised transfer learning for robust, scalable, and interpretable integration of single-cell multi-omics data. Code is available at https://github.com/No-AgCl/GuidedCoC.

## 1 Introduction

Recent advances in single-cell multi-omics technologies have revolutionized our ability to dissect cellular heterogeneity by simultaneously profiling gene expression (scRNA-seq) and chromatin accessibility (scATAC-seq) at single-cell resolution [1]. These paired modalities offer complementary insights: scRNA-seq captures transcriptional states, while scATAC-seq identifies regulatory elements governing cell identity and function [2, 3]. However, joint analysis of these data remains challenging due to technical noise, sparsity, and batch effects, particularly in emerging datasets where paired measurements are often limited in quality or scale [4]. Concurrently, large-scale single-cell atlases, such as Tabula Muris and Human Cell Atlas, provide high-quality unpaired scRNA-seq data with inherent cluster structures that approximate known cell types [5, 6]. This presents an opportunity to leverage such biologically meaningful structures to enhance analysis of sparse, noisy multi-omics data - a paradigm shift from traditional single-dataset clustering.

Current computational strategies for multi-omics integration, including matrix factorization (MOFA+) and graph-based alignment (Seurat v4), primarily focus on harmonizing paired modalities [7, 8]. While effective, these methods often overlook the potential of external, unpaired datasets to scaffold clustering in target data. Furthermore, they struggle with feature-level alignment, such as linking gene modules in scRNA-seq to regulatory peaks in scATAC-seq, which is critical for interpreting transcriptional regulation [9]. Transfer learning approaches in single-cell analysis have shown promise in propagating annotations across datasets [10, 11], but they typically require explicit labels and fail to exploit shared feature spaces across modalities.

Here, we address these gaps by introducing Guided Co-clustering Transfer (GuidedCoC), a framework that integrates paired scRNA-seq/scATAC-seq target data (ℛ^(*t*)^, 𝒜^(*t*)^) with unpaired source scRNA-seq (ℛ^(*s*)^) through information-theoretic co-clustering. Our approach is biologically motivated by two observations: (1) High-quality scRNA-seq datasets, even without explicit labels, encode robust cluster structures reflective of cell identity [5]; (2) Gene activity score or promoter accessibility in scATAC-seq provide a natural bridge to scRNA-seq features, enabling cross-modal feature co-clustering [12]. By formulating a joint optimization over cell and feature clusters, GuidedCoC transfers knowledge from ℛ^(*s*)^ to stabilize clustering in ℛ^(*t*)^/ 𝒜^(*t*)^, while aligning gene-peak modules to enhance interpretability.

### Our Contributions

This work makes the following key contributions: **(1) New Problem Setting**. We investigate a realistic yet underexplored setting: knowledge transfer across unpaired and paired single-cell multi-omics data. Specifically, we leverage unsupervised cluster structures in high-quality, unpaired scRNA-seq source data to guide clustering in paired scRNA-seq/scATAC-seq target data, thereby avoiding reliance on explicit annotations. **(2) Novel Methodology**. We propose GuidedCoC, an improved information-theoretic co-clustering framework [13], to simultaneously (a) co-cluster gene expression and chromatin accessibility features - aligning transcriptional modules with regulatory elements - and (b) cluster cells in paired multi-omics data. Cross-modal feature co-clustering implicitly performs dimension reduction in a shared feature space, mitigating noise and sparsity, and enhancing downstream cell clustering. GuidedCoC also automatically aligns similar cell types across unpaired and paired data, bridging the gap between single-modality transfer learning and multi-omics integration. **(3) Comprehensive Evaluation**. Extensive experiments on multiple benchmark datasets demonstrate the superiority of GuidedCoC over state-of-the-art methods in both clustering accuracy and biological interpretability. Our results highlight GuidedCoC’s potential for scalable and robust analysis of noisy, sparse single-cell multi-omics data.

## 2 Related Work

Integrating single-cell multi-omics data and transferring biological structure across datasets are active and complementary research areas. We organize our discussion into two key directions:

### Single-Cell Multi-Omics Integration

Early efforts in integrating paired scRNA-seq and scATAC-seq data primarily focused on latent space alignment across modalities. Methods such as MOFA+ [7] and Seurat v4 [8] apply dimensionality reduction to jointly embed multi-modal data, facilitating downstream clustering and visualization, but they often neglect feature-level alignment and modality-specific signal. More expressive models, such as Cobolt [14], leverage hierarchical variational autoencoders to disentangle shared and private representations, while BABEL [15] performs cross-modal prediction to impute missing modalities. MultiVI [16] further extends probabilistic modeling to integrate and augment sparse, paired multi-omics profiles. However, these methods largely rely on fully paired data and can struggle under extreme sparsity [17]. To move beyond full pairing, scMoMaT [18] addresses mosaic integration through matrix tri-factorization, aligning cells and features across batches and modalities. Nevertheless, it does not directly target the clustering of paired data or incorporate external guidance. A recent benchmarking study [19] assesses many of these methods, yet it overlooks the potential of external information to improve clustering and interpretability in paired multi-omics contexts. Another underdeveloped aspect is cross-modal feature alignment. Peak-to-gene linkage methods such as Cicero [9] and MAESTRO [20] statistically correlate chromatin accessibility peaks with gene expression, but these tools operate outside a unified co-clustering framework and do not leverage external reference data. In contrast, our method explicitly performs joint co-clustering of genes and regulatory features across modalities, and uniquely incorporates unpaired scRNA-seq data to guide integration enabling more interpretable and stable results in the presence of noise and sparsity.

### Transfer Learning in Single-Cell Analysis

Transfer learning has seen increasing application in single-cell biology, particularly for label propagation and domain adaptation. Approaches such as scArches [21] utilize conditional variational autoencoders to map new datasets onto reference atlases, while singleCellNet [22] transfers cell-type annotations using ensemble-based classifiers. These methods, however, require explicit source labels and are primarily designed for single-modality data. Unsupervised transfer learning remains less explored in this domain. SCOT [23] applies optimal transport to align datasets without supervision, but focuses on cell-level correspondences and overlooks feature relationships. SMAI [24] achieves structure-preserving alignment across datasets but assumes shared cell types and does not operate in the multi-omics setting. Our work diverges from these paradigms by transferring latent feature clusters - derived from unpaired scRNA-seq datasets - to improve co-clustering in paired multi-omics data. This novel strategy is motivated by the biological insight that transcriptional programs (e.g., gene modules regulated by lineage-specific transcription factors) are often conserved and transferable across conditions [5, 25]. Large-scale efforts such as Tabula Muris [5] and the Human Cell Atlas [6] show that unsupervised clustering in scRNA-seq can approximate canonical cell types, underscoring the value of structure-guided transfer even without explicit labels. Recent reviews [26] have highlighted the gap between unsupervised biology and interpretable, integrative analysis - our work seeks to bridge this gap through guided co-clustering and feature alignment informed by cross-dataset regulatory logic.

## 3 The Methodology

### 3.1 Problem Setting

We address an unsupervised transfer clustering problem involving three datasets: (i) paired scRNA-seq data ℛ^(*t*)^ and scATAC-seq data 𝒜^(*t*)^, which constitute the *target* and are typically noisy and sparse; and (ii) unpaired scRNA-seq data ℛ^(*s*)^, serving as the *source*, assumed to be of higher quality with robust cluster structure but without explicit labels. We assume that a subset of features in 𝒜^(*t*)^ - specifically gene activity score or promoter accessibility - are linked to ℛ^(*t*)^ and share clustering structure with the source scRNA-seq data ℛ^(*s*)^. This enables potential alignment at both the cell and feature levels. Our goal is to transfer structural knowledge - cell clusters and feature modules - from the source data to improve clustering and feature alignment in the target multi-omics data. This setting captures a biologically realistic scenario: unpaired scRNA-seq datasets often contain high-quality unsupervised clusters (e.g., Tabula Muris [5]), which reflect conserved gene modules and regulatory programs. We aim to exploit this structure to guide co-clustering in sparse paired data. To this end, we propose a unified information-theoretic framework that jointly co-clusters cells and features across modalities and domains, enabling cross-modal integration, noise reduction, and biologically meaningful clustering without requiring annotations.

### 3.2 Co-clustering for Each Dataset

We begin by considering the target data ℛ^(*t*)^, represented as an *n*^(*t*)^ × *q* matrix, containing *q* features measured across *n*^(*t*)^ cells. Let *Y* and *Z* denote discrete random variables corresponding to cell and feature indices, respectively, where *Y* ∈ {1, …, *n*^(*t*)^} and *Z* ∈ {1, …, *q*}. The joint probability 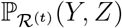 captures the activity of feature *z* in cell *y*, estimated by normalizing the data matrix so that the total sum equals one:

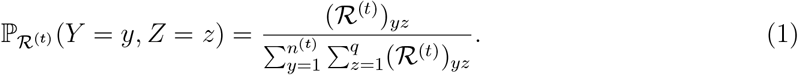

The goal of co-clustering is to group similar cells and features. Assume *N* ^(*t*)^ cell clusters and *K* feature clusters. Define 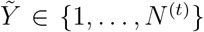 and 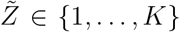 as the corresponding cluster variables, determined by clustering functions *C*_*Y*_ and *C*_*Z*_, such that 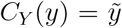 and 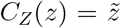. The joint distribution over clusters is then:

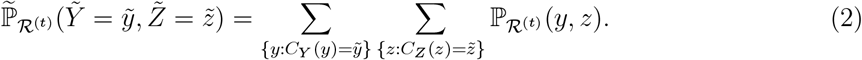

For the target domain’s paired dataset ℛ^(*t*)^, we define a clustering loss grounded in the KL divergence between the joint cell-feature distribution and its constructed reference:

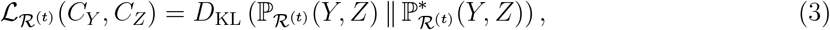

where the reference 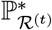 adjusts a coarse-grained distribution using marginals to approximate cluster-based independence, and 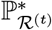 is structured as:

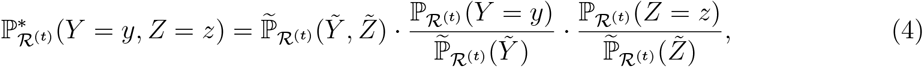

where 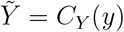 and 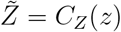 are the cluster indices. The objective 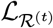 in Eq. (3) equivalently minimizes the loss of mutual information between cells and features during co-clustering, as established in the information-theoretic co-clustering framework [13, 27].

For the auxiliary target modality 𝒜^(*t*)^, represented as an *n*^(*t*)^ × *q* matrix, sharing cell and feature identities with ℛ^(*t*)^ (features from both datasets are profiled from the same single cell and are linked), we reuse *Y*, *C*_*Y*_, *Z* and *C*_*Z*_:

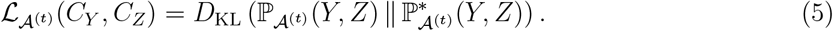

For the source data ℛ^(*s*)^ (where the cell clustering function *C*_*X*_ is known with |*C*_*X*_| = *N* ^(*s*)^ clusters, and feature identities are shared across domains), represented as an *n*^(*s*)^ × *q* matrix, we reuse *Z* and *C*_*Z*_:

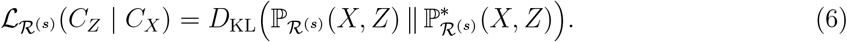

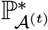 and 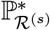 mirror 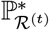‘s structure (Eq. (4)).

### 3.3 Guided Co-clustering Transfer Learning

We integrate paired/unpaired single-cell multi-modal data via joint clustering of cells and features across domains. The framework has two stages: (1) unsupervised co-clustering of ℛ^(*t*)^, 𝒜^(*t*)^, and ℛ^(*s*)^, and (2) cross-domain matching of shared cell populations.

#### 3.3.1 Stage 1: Integrative Co-Clustering

The final co-clustering objective jointly minimizes loss across modalities and domains:

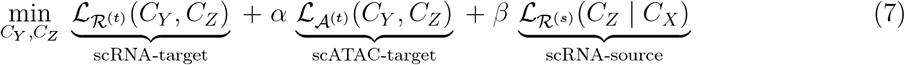

with hyperparameters *α, β* controlling modality and domain contributions. The key idea is that the clustering function *C*_*Z*_ over features acts as a conduit for transferring knowledge from the source data to the target domain. Given high-quality clustering *C*_*X*_ in ℛ^(*s*)^, the learned feature clusters *C*_*Z*_ improve both integration and robustness across domains. Furthermore, by coupling ℛ^(*t*)^ and 𝒜^(*t*)^ via shared clustering functions, we leverage modality-specific structure to enhance biological interpretability. Here, we empirically set the number of shared feature clusters to *K* = 12, and the weighting hyperparameters to *α* = 0.8 and *β* = 1. For a detailed sensitivity analysis, please refer to Figure 4 in Section 4.2.

##### Optimization

The KL divergence for ℛ^(*t*)^ (Eq. (3)) admits two symmetric decompositions - conditioning on cells or features - highlighting local consistency of cluster assignments [27, 28]:

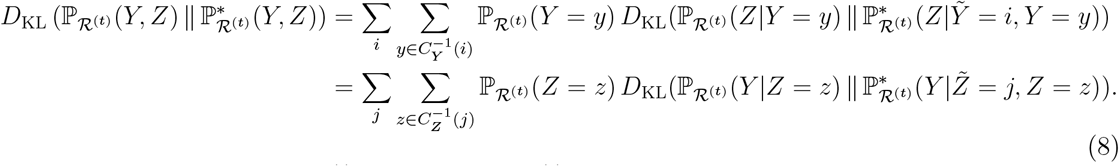

The KL divergences for 𝒜^(*t*)^ (Eq. (5)) and ℛ^(*s*)^ (Eq. (6)) also mirror this structure. We optimize Eq. (7) via block-coordinate descent:

- Updating *C*_*Y*_ (cell clusters): Assign each *y* to cluster *i* via minimizing the weighted sum of KL divergences between conditional feature distributions in ℛ^(*t*)^ and 𝒜^(*t*)^(Eq. (A2)).
- Updating *C*_*Z*_ (feature clusters): Assign each *z* to cluster *j* via minimizing a sum of KL divergences across all datasets: ℛ ^(*t*)^, 𝒜^(*t*)^, and ℛ ^(*s*)^ (Eq. (A3)).

The *C*_*Y*_ update ensures cell-cluster consistency across real and augmented data; the *C*_*Z*_ update identifies functionally coherent features preserved across domains. Iterative refinement yields a unified latent representation, converging within 10 iterations (Figure 5). Details (derivations, algorithm, complexity, convergence) are in Appendix A.

#### 3.3.2 Stage 2: Cross-Domain Cluster Matching

Given learned target domain clusters *C*_*Y*_ and known source clusters *C*_*X*_, the goal of the second stage is to align clusters that represent shared biological cell types. We construct cluster-level joint distributions over features, denoted 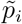 for source and 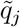 for target clusters. Matching is performed by minimizing the average Jensen-Shannon divergence (AJSD) across *n*_trials_ random subsamples:

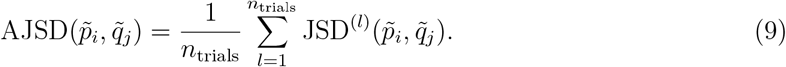

Only cluster pairs with AJSD below a threshold *τ*_JSD_ are retained as valid matches. To avoid assignment conflicts and ensure optimal alignment, we repeat the matching over *n*_shuffles_ permutations of *C*_*Y*_, selecting the configuration ℳ* that minimizes the summation of AJSD over all valid matches:

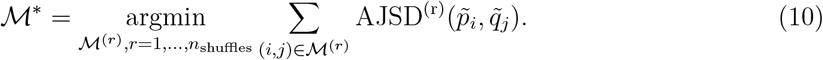

Random permutations help resolve suboptimal greedy assignments and enforce exclusivity (each source cluster can be matched once per shuffle). This enhances robustness and improves biological plausibility by selecting configurations with clearer inter-cluster separation. We set *n*_trials_ = *n*_shuffles_ = 25, *τ*_JSD_ = 0.45. Details (derivations, algorithm, complexity) are in in Appendix B.

We name our framework **GuidedCoC** to highlight its core mechanism: a *Guided Co-clustering* strategy in which structural information from high-quality unpaired scRNA-seq data informs both the joint co-clustering of cells and features in paired scRNA-seq/scATAC-seq data, and the cross-domain cluster alignment of shared cell populations; see Figure 1 for an illustrative toy example.

**Figure 1:**
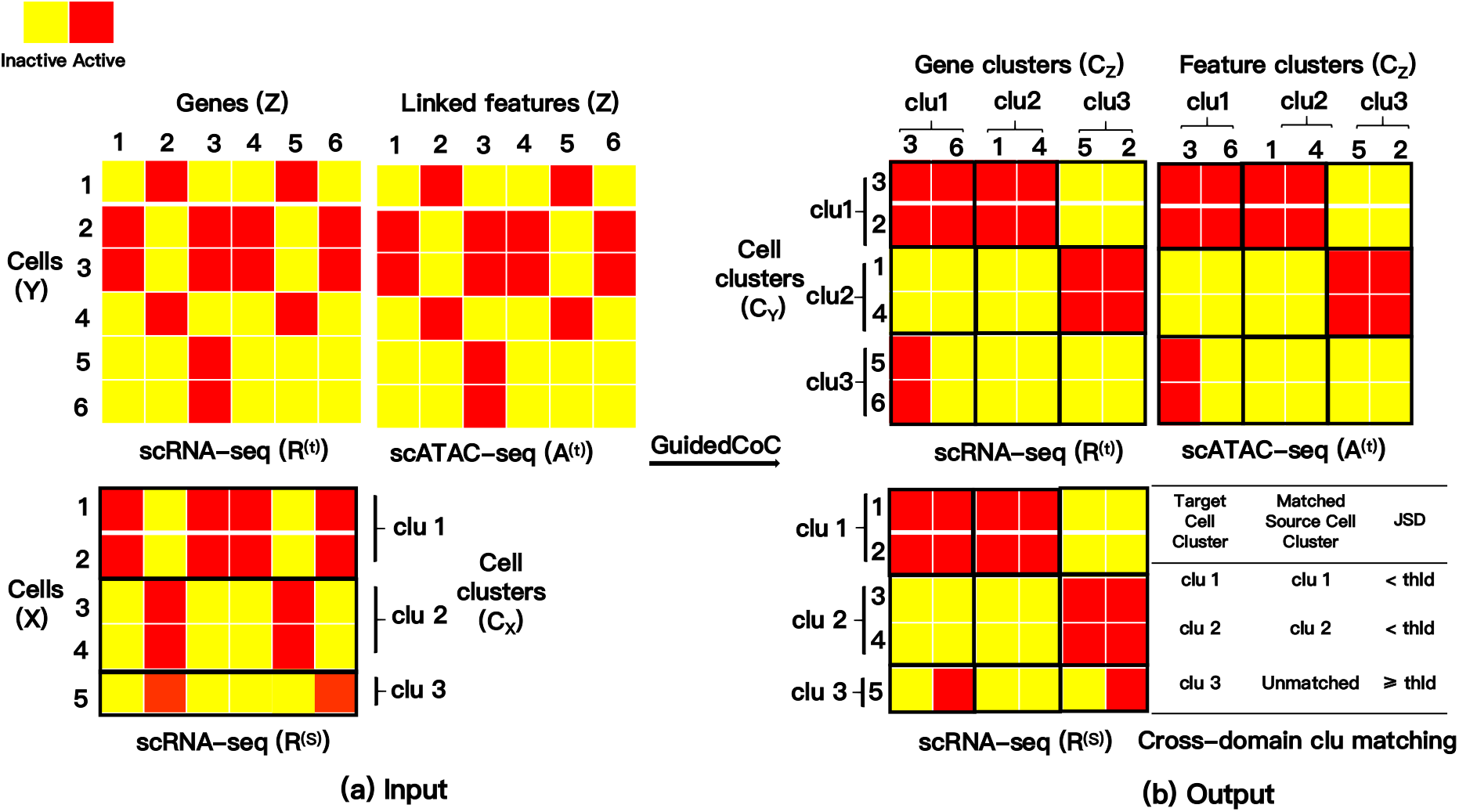
A toy example of **GuidedCoC**. (a) The input includes paired target datasets ℛ^(*t*)^ and 𝒜^(*t*)^, where gene activity scores (or promoter accessibility) in 𝒜^(*t*)^ are linked to genes in ℛ^(*t*)^, and a source scRNA-seq dataset ℛ^(*s*)^ that shares the same gene set and has known cell clusters *C*_*X*_. (b) The output includes shared feature clusters *C*_*Z*_, target cell clusters *C*_*Y*_, and cross-domain cluster matching for shared cell populations between the source and target data. “clu”: cluster; “thld”: threshold.

### 3.4 Feature Selection and Data Preprocessing

We begin by selecting highly variable genes (HVGs) from the source scRNA-seq dataset ℛ^(*s*)^ using Seurat [29]. To ensure consistency in the feature space, the same set of HVGs is used for the target scRNA-seq dataset ℛ^(*t*)^, with homologous genes selected in the case of cross-species integration. Empirically, a selection of 2,500 HVGs is sufficient to capture the key transcriptional variability across cells. Both source and target scRNA-seq datasets are normalized using a log_2_(· + 1) transformation, applied to UMI counts or TPM values depending on the data type. For the scATAC-seq modality 𝒜^(*t*)^, we compute gene activity scores by counting the number of fragments overlapping the gene body and a 2-kb upstream promoter region - following the implementation of the `GeneActivity` function in the `Signac` package [30]. These scores are then log-transformed using log_2_(· + 1) prior to model input to stabilize variance. Importantly, GuidedCoC automatically adjusts for sequencing depth during training, thereby obviating the need for explicit normalization across modalities.

## 4 Experiments

### 4.1 Experimental Settings

#### Evaluation Metrics

We assess clustering performance on the target data (ℛ^(*t*)^) using two standard metrics: Normalized Mutual Information (NMI) and Adjusted Rand Index (ARI) [31] (definitions in Appendix D). Higher values of both metrics indicate better cluster agreement with ground truth.

#### Datasets

We evaluate on four real-world multiome benchmarks spanning diverse tissues (PBMC, embryonic mouse brain, lymph node/spleen, and pancreatic islets), species (human and mouse), and sequencing platforms, all publicly available with details in Appendix E and Table 1. Both source (ℛ^(*s*)^) and target (ℛ^(*t*)^) datasets provide known cell clusters (*C*_*X*_ for ℛ^(*s*)^ and ground truth for ℛ^(*t*)^ evaluation). For Example 1, we leverage the well-annotated ℛ^(*s*)^ to predict ℛ^(*t*)^ cell types through cross-domain matching.

**Table 1:** Statistics of unpaired and paired multi-omic single-cell datasets used in our experiments.”PBMC”: Peripheral Blood Mononuclear Cells; “MB”: Mouse Brain; “LN”: Lymph Node; “PI”: Pancreatic Islet.

| Datasets | Source Data<br>($\mathcal{R}^{(s)}$) | #Cells<br>($n^{(s)}$) | #Cell Types<br>($N^{(s)}$) | Target Data<br>($\mathcal{R}^{(t)}/\mathcal{A}^{(t)}$) | #Cells<br>($n^{(t)}$) | #Cell Types<br>($N^{(t)}$) |
| --- | --- | --- | --- | --- | --- | --- |
| Example1 | scRNA-seq (PBMC, annotated) | 2,283 | 7 | Multiome (PBMC) | 3,010 | 8 |
| Example2 | scRNA-seq (E18 MB) | 2,247 | 12 | Multiome (E18 MB) | 3,276 | 9 |
| Example3 | scRNA-seq (Mouse LN) | 2,555 | 11 | Multiome (Human LN) | 2,426 | 13 |
| Example4 | scRNA-seq (Human PI) | 6,371 | 14 | Multiome (Human PI) | 22,146 | 8 |

**Table 2:** Quantitative comparisons with state-of-the-art approaches across four real-world examples. **Bold** indicate best results. <u>Underline</u> indicate second-best results.

| Methods | Example 1 |  | Example 2 |  | Example 3 |  | Example 4 |  |
| --- | --- | --- | --- | --- | --- | --- | --- | --- |
|  | NMI | ARI | NMI | ARI | NMI | ARI | NMI | ARI |
| GuidedCoC | **0.6676** | **0.5399** | **0.5640** | **0.4089** | **0.5558** | **0.4539** | **0.7570** | **0.8224** |
| MultiVI | 0.6326 | <u>0.5259</u> | <u>0.4928</u> | 0.3240 | <u>0.4823</u> | <u>0.3542</u> | 0.5235 | 0.3131 |
| Cobolt | 0.5420 | 0.4090 | 0.4785 | <u>0.3392</u> | 0.4319 | 0.2428 | <u>0.5847</u> | <u>0.5633</u> |
| ScMoMaT | 0.5259 | 0.4068 | 0.4751 | 0.3269 | 0.4256 | 0.2922 | 0.2572 | 0.1162 |
| Seurat v4 | <u>0.6376</u> | 0.4764 | 0.4845 | 0.3148 | 0.4726 | 0.2600 | 0.3777 | 0.1472 |

#### Benchmark Comparisons

We evaluate against four state-of-the-art baselines spanning different approaches: MultiVI [16] (deep generative), Cobolt [14] (generative), scMoMaT [18] (matrix factorization), and Seurat v4 [8] (graph-based). All methods use their standard preprocessing pipelines, with cell clusters obtained via Leiden algorithm [32] using ground-truth cluster numbers (*N* ^(*t*)^) from the original datasets to ensure fair comparison (consistent with our GuidedCoC implementation). Implementation details follow [19], with full method descriptions in Appendix F.

### 4.2 Experimental Results

#### Performance Comparison

To evaluate clustering performance on the target scRNA-seq data ℛ^(*t*)^, we conduct extensive experiments comparing GuidedCoC with state-of-the-art baselines (Ta-ble 2). Our method demonstrates consistent improvements in NMI and ARI across all benchmarks, particularly in Example 4, attributable to two key advantages: (1) explicit utilization of known cell-population clusters *C*_*X*_ from unpaired source data ℛ^(*s*)^ (where high-quality cluster structures enable effective knowledge transfer), and (2) unified feature alignment across ℛ^(*s*)^, ℛ^(*t*)^, and 𝒜^(*t*)^ via shared clustering function *C*_*Z*_, which jointly addresses cross-domain and cross-modality integration challenges. These synergistic mechanisms collectively explain GuidedCoC’s superior performance in both quantitative metrics and visual assessments (Figure 2).

**Figure 2:**
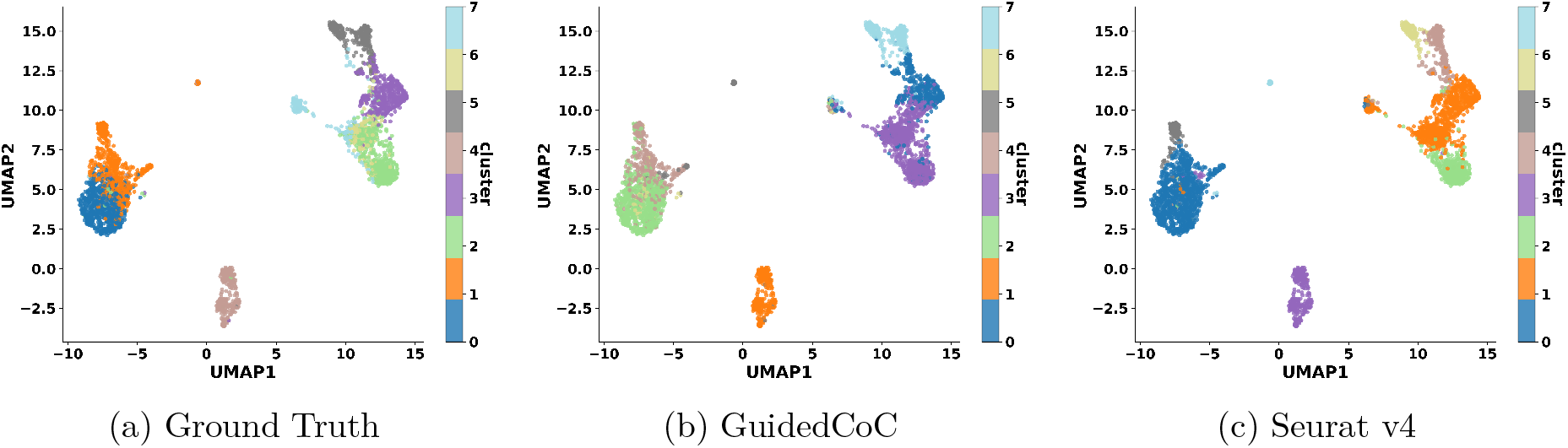
UMAP visualization of cells in the target scRNA-seq dataset ℛ^(*t*)^ (Example 1).

#### UMAP Visualization

Figure 2 presents a UMAP visualization of the target scRNA-seq data ℛ^(*t*)^ from Example 1, comparing clustering results from GuidedCoC, Seurat v4, and the ground-truth annotations. GuidedCoC successfully captures the major cell populations with improved alignment to the ground truth relative to Seurat v4. Notably, our method preserves the separation between biologically distinct yet spatially adjacent clusters - such as clusters 0 and 1, and clusters 2 and 3 from ground truth annotations - that Seurat v4 tends to partially merge. These results highlight the strength of GuidedCoC in preserving both global separability and local structural consistency during multi-modal integration. More visualization results are available at Appendix G.

#### Ablation Study

We conduct a comprehensive ablation study to evaluate the contributions of key components in GuidedCoC (Table 3). Two variants are analyzed: (1) *GuidedCoC w/o* ℛ^(*s*)^, which excludes source data by setting *β* = 0 in Eq. (7), and (2) *GuidedCoC w/o* 𝒜^(*t*)^, which omits auxiliary paired data by setting *α* = 0 in Eq. (7). The observed performance degradation in both variants - measured by NMI and ARI - underscores the critical role of external knowledge transfer (via ℛ^(*s*)^) and cross-modality integration (via 𝒜^(*t*)^) for robust multi-omics analysis. This validates our design choices for joint feature and sample alignment.

**Table 3:** Ablation study of GuidedCoC across four examples. **Bold** indicate best results.

|  | Example 1 |  | Example 2 |  | Example 3 |  | Example 4 |  |
| --- | --- | --- | --- | --- | --- | --- | --- | --- |
|  | NMI | ARI | NMI | ARI | NMI | ARI | NMI | ARI |
| GuidedCoC | **0.6676** | **0.5399** | **0.5640** | **0.4089** | **0.5558** | **0.4539** | **0.7570** | **0.8224** |
| GuidedCoC w/o $\mathcal{R}^{(s)}$ | 0.6478 | 0.4790 | 0.5580 | 0.3948 | 0.5508 | 0.4431 | 0.6827 | 0.7605 |
| GuidedCoC w/o $\mathcal{A}^{(t)}$ | 0.6416 | 0.4679 | 0.5274 | 0.3696 | 0.5382 | 0.3609 | 0.7232 | 0.5375 |

#### Biological Interpretation

We examine the biological relevance of feature clusters identified by GuidedCoC in Example 1 (Figure 3), focusing on the three best-matched cell-type pairs based on their top-three Jensen-Shannon divergence (JSD) scores (right panel of Figure 3). Feature clusters clu1 and clu2 are predominantly associated with cell cluster `clu6` in the target scRNA-seq dataset ℛ^(*t*)^, which our method matched to T cells in the source dataset ℛ^(*s*)^ through cross-domain cell cluster alignment. Additionally, feature clusters `clu7–clu9` are enriched in cell clusters `clu2` and `clu4`, corresponding to lymphocytes and B cells in ℛ^(*s*)^, respectively. To elucidate the biological functions of these feature clusters, we perform functional enrichment analysis using DAVID [33, 34]. Genes in `clu1` show significant enrichment for terms such as *T cell receptor complex* (Bonferroni-corrected *p*-value = 5.68 × 10^−5^), reflecting the core structural and signaling components of the TCR. Genes in `clu2` are strongly associated with *antigen processing and presentation* and *positive regulation of T cell proliferation* (Bonferroni-corrected *p*-values = 4.51 × 10^−9^ and 1.07 × 10^−6^, respectively), underscoring their role in T cell activation and immune checkpoint regulation. For full details, see Table A1 in Appendix H. Similarly, genes in `clu7–clu9` are linked to receptor-mediated activation, cell adhesion, and cytotoxic functions in B cells and lymphocytes (see Table A2 in Appendix H). These results validate the interpretability of our framework and demonstrate its ability to uncover biologically meaningful relationships between cell types and feature clusters.

**Figure 3:**
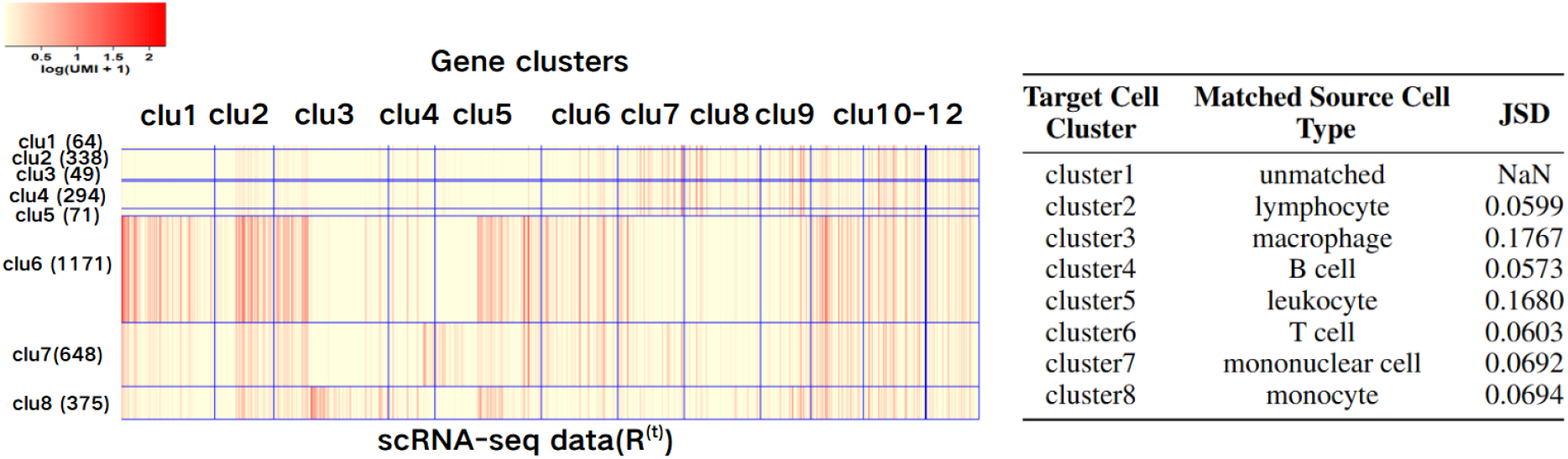
Biological interpretation of GuidedCoC (Example 1). **Left**: Heatmap of the clustering results for ℛ^(*t*)^ (labeled by “clu” for cluster). For clarity, pseudocells are generated by randomly averaging every 20 cells within the same cluster. **Right**: Cross-domain cell cluster matching results from stage 2 of GuidedCoC. “NaN” reflects full alignment of all seven source cell types to target clusters, with no residual unmatched populations.

#### Sensitivity Analysis

We evaluate the robustness of GuidedCoC with respect to three key hyperparameters: the number of feature clusters *K*, the modality weighting factor *α* for the scATAC-seq-derived gene activity/promoter accessibility matrix 𝒜^(*t*)^, and the domain weighting factor *β* for the source scRNA-seq data ℛ^(*s*)^. Figure 4 reports NMI scores across all four examples under varying hyperparameter values, with others held fixed: (1) **Feature cluster count (***K*): Panel (a) shows performance for *K* ∈ {2, 4, …, 64}. NMI improves monotonically until plateauing at *K* = 12, suggesting this value sufficiently captures both modality- and domain-shared structure. We fix *K* = 12 in all experiments. (2) **Modality weight (***α*): Panel (b) demonstrates stable

**Figure 4:**
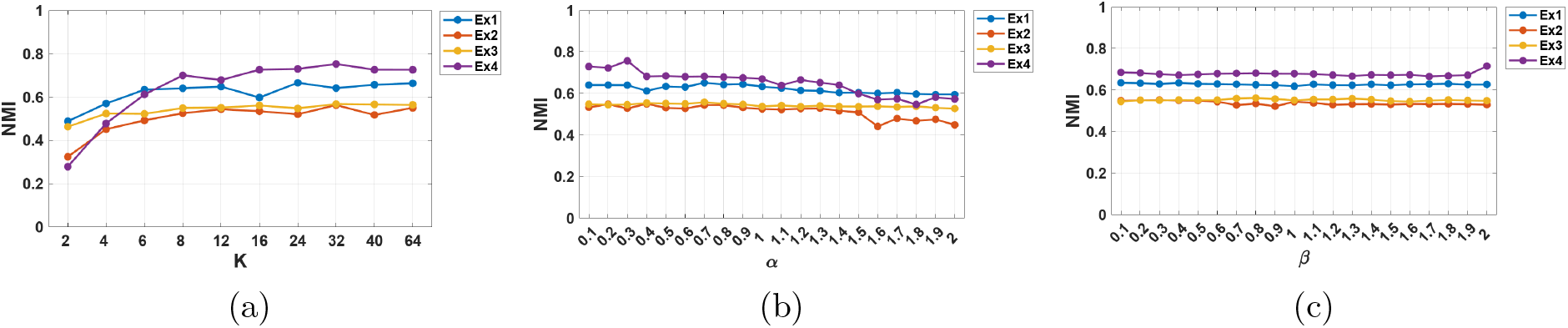
Sensitivity to the number of feature clusters *K* and two weighting factors *α, β* across four examples.

NMI for *α* ∈ (0, 1.3], with degradation outside this range. The decline at high *α* reflects noise in scATAC-seq data; we empirically set *α* = 0.8 to balance information from 𝒜^(*t*)^. (3) **Domain weight (***β*): Performance remains robust across *β* ∈ (0, 2] (Panel (c)), indicating insensitivity to source domain weighting. We use *β* = 1 as default. GuidedCoC exhibits consistent performance across wide hyperparameter ranges, demonstrating its reliability for multi-modal, cross-domain single-cell integration.

#### Convergence and Efficiency

The left panel of Figure 5 illustrates the convergence behavior of the normalized objective value over iterations. Across all four examples, the objective decreases rapidly during the initial iterations and stabilizes thereafter, demonstrating both the efficiency and stability of the optimization procedure in GuidedCoC. The right panel of Figure 5 compares the runtime performance of GuidedCoC with baseline methods. Our method exhibits competitive computational efficiency, particularly when clustering large-scale datasets with approximately 30,000 cells (e.g., Example 4), completing in under 10 minutes and ranking second overall. We attribute this favorable runtime to the use of parallel computing in both stages of our framework (see Appendix C for details on the parallelization strategy).

**Figure 5:**
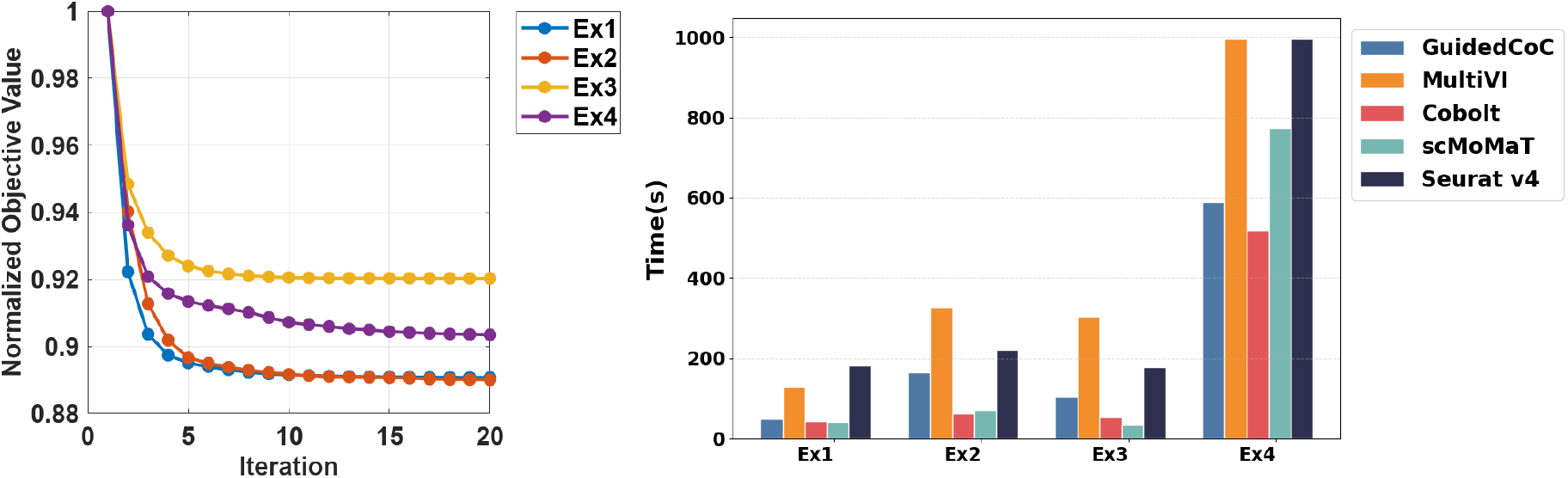
Convergence and computational performance. **Left**: Loss function convergence curves for GuidedCoC. **Right**: Running time comparison across datasets.

## 5 Discussion

We have presented **Guided Co-clustering Transfer (GuidedCoC)**, an unsupervised framework for integrative analysis of paired single-cell RNA-seq (scRNA-seq) and ATAC-seq (scATAC-seq) data guided by unpaired reference scRNA-seq datasets, addressing the critical challenge of cross-modal knowledge transfer in single-cell multi-omics analysis. By jointly co-clustering cells and features across modalities and domains through an information-theoretic objective, GuidedCoC simultaneously (1) aligns gene expression and chromatin accessibility patterns, (2) enhances cell-type resolution via cross-modal regularization, and (3) automatically identifies shared cell populations between source and target datasets. Empirical results demonstrate that GuidedCoC consistently outperforms state-of-the-art methods in clustering accuracy while achieving superior biological interpretability and computational scalability, establishing unsupervised structural transfer as a powerful approach for robust single-cell data integration.

### Broader Impact

Our framework offers a scalable and annotation-free approach to multi-modal data integration, particularly valuable when high-quality paired measurements are limited. GuidedCoC can empower downstream biological discovery by improving cell-type resolution and uncovering interpretable feature modules linked across omics layers. More broadly, the idea of leveraging structural priors from large, unpaired datasets opens new opportunities for integrating diverse data modalities in other domains such as spatial transcriptomics or single-cell proteogenomics. As single-cell atlases continue to grow, methods like GuidedCoC that leverage their structure without requiring explicit labels are increasingly impactful.

### Limitations and Future Work

Despite its promising performance, GuidedCoC assumes the availability of a high-quality source dataset and partial overlap between cell populations in source and target datasets. In scenarios where the source and target domains are highly divergent, performance may degrade due to distributional mismatch. Moreover, the current framework focuses on gene activity scores/promoter accessibility as the links between modalities; extending it to accommodate more complex regulatory relationships (e.g., enhancer-promoter interactions or TF binding motifs) is an important direction for future research. Finally, formalizing theoretical guarantees for transferability and extending the approach to support more than two omics layers would further enhance its utility in comprehensive single-cell multi-omics integration.

## A Optimization for Integrative Co-clustering

### A.1 Mathematical Derivation

The goal is to jointly infer the cluster assignments *C*_*Y*_ (target domain cell clusters) and *C*_*Z*_ (feature clusters) by minimizing the divergence between the observed joint distributions and their cluster-induced approximations. Specifically, we solve the following optimization problem:

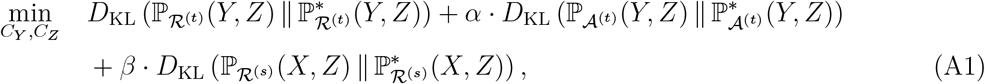

where *α* and *β* are tunable trade-off parameters. Each KL divergence term quantifies how well the learned clustering structure captures the joint dependency of variables in different data modalities or domains.

To minimize this objective, we adopt an alternating optimization strategy by iteratively updating *C*_*Y*_ and *C*_*Z*_, while keeping the other fixed.

1. **Fix** *C*_*X*_: The cell type labels in the source domain ℛ^(*s*)^ are assumed known and fixed, so *C*_*X*_ is not updated during optimization.
2. **Update** *C*_*Y*_ **given** *C*_*Z*_: With feature clusters *C*_*Z*_ fixed, we update the cell clustering in the target domain by minimizing the following objective:

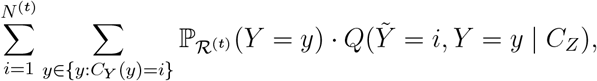

where the per-sample cost function 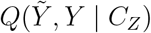 is defined as:

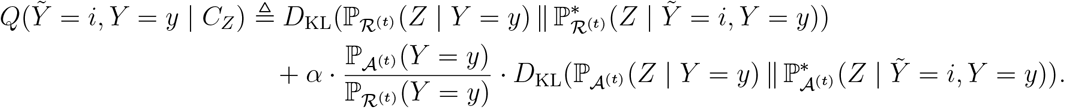

Each cell *y* ∈ ℛ^(*t*)^ is assigned to the cluster *i* that minimizes the local cost:

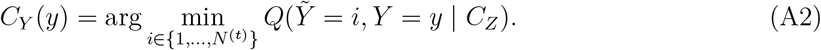
3. **Update** *C*_*Z*_ **given** *C*_*Y*_ : With the target cell clusters fixed, we next update the feature clusters by minimizing:

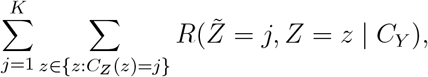

where the function 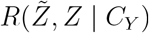 aggregates divergence costs across domains:

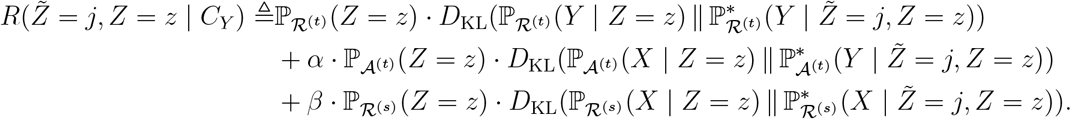

Each feature *z* ∈ {1, …, *q*} is assigned to the cluster *s* that minimizes the above expression:

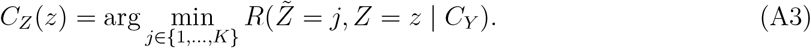

### A.2 Integrative Co-Clustering Algorithm

We provide the detailed algorithm of the optimization for integrative co-clustering in Algorithm 1.

#### Algorithm 1 Integrative Co-Clustering

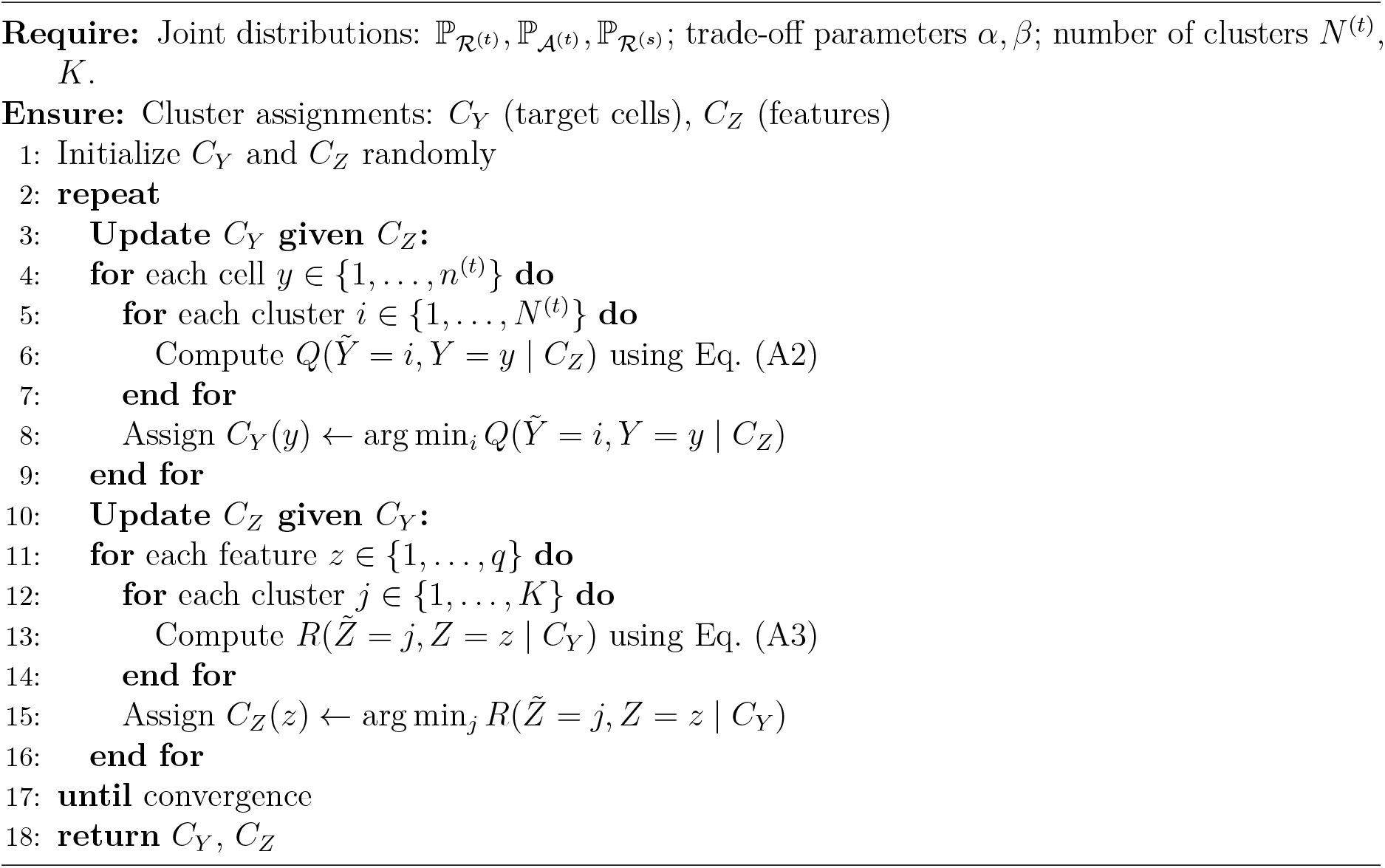

#### Complexity of Algorithm 1

Here we consider per-iteration cost. Let *n*^(*t*)^ = #*row of* ℛ^(*t*)^, *n*^(*s*)^ = #*row of* ℛ^(*s*)^, *q* = #*features, N* ^(*t*)^ = #*cell* − *clusters*, and *K* = #*feature* − *clusters*.

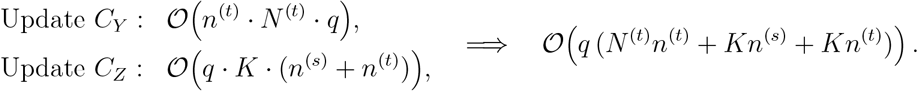

The calculation of complexity has ignored the multiples of constants, and the complexity of calculating 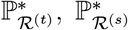 and 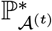 has been omitted (in the process of calculating *Q*, and the complexity is 𝒪((*n*^(*s*)^ + *n*^(*t*)^)*q*), so it is not included in the total complexity).

### A.3 Convergence

The following theorem shows that the Algorithm 1 converges to a coordinate-wise stationary point. It indicates the algorithm will converge to a local minimum. Finding the global optimal solution is NP-hard.

#### Theorem A.1.

*Let* 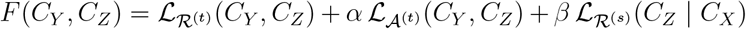 *and assume each divergence is finite. Then the alternating updates*

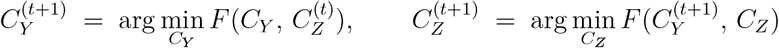

*generate a sequence* 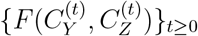 *that is* monotone nonincreasing, lower-bounded *by zero, and converges to a limit value F*\*. *Moreover, any limit point* 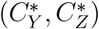 *of the sequence of assignments satisfies the coordinate-wise optimality conditions*

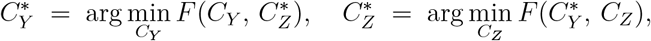

*i*.*e. is a* coordinate-wise stationary point *of F*.

*Proof*. We break the proof into three parts.

#### (a) Monotonicity

At iteration *t*, first update

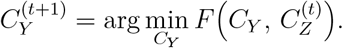

By choosing the global minimizer in *C*_*Y*_ (over a finite discrete set) we have

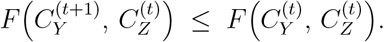

Next, with *C*_*Y*_ fixed, update

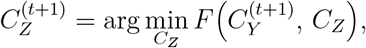

which similarly yields

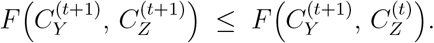

Combining the two inequalities establishes

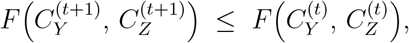

so the sequence 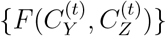 is monotone nonincreasing.

#### (b) Lower-Boundedness

Each KL divergence *D*_KL_(·∥·) is nonnegative and finite by assumption, hence *F* (*C*_*Y*_, *C*_*Z*_) ≥ 0 for all assignments. Thus the monotone nonincreasing sequence is bounded below by zero.

#### (c) Convergence and Stationarity

A real sequence that is monotone nonincreasing and bounded below converges to a finite limit *F*\*. Denote by 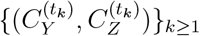 a subsequence converging to some limit point 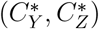 (which exists because the assignment sets are finite).

We now show that 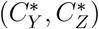 satisfies the coordinate-wise optimality conditions. By continuity of *F* in its discrete arguments (i.e. it takes only finitely many values), the limiting arguments satisfy

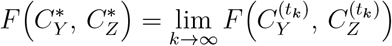

and, since each 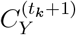 minimized 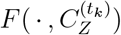,

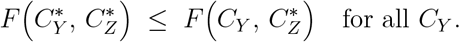

Similarly, from the minimization in the *C*_*Z*_-update,

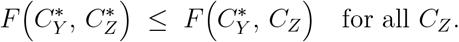

These two inequalities jointly imply that 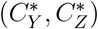 is a coordinate-wise stationary point:

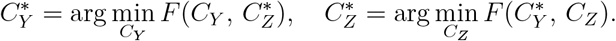

This concludes the proof. □

## B Details for Cross-Domain Cluster Matching

### B.1 Derivation

Let *C*_*X*_ denote the known cell type labels in the source domain ℛ^(*s*)^, and let *C*_*Y*_ represent the clustering results inferred in the target domain ℛ^(*t*)^. We assume that a subset of clusters in *C*_*X*_ and *C*_*Y*_ correspond to shared biological cell types across domains. Our goal is to establish one-to-one correspondences between such clusters, enabling accurate label transfer from ℛ^(*s*)^ to ℛ^(*t*)^.

Let 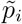 and 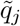 denote the cluster-level feature profiles derived from the *i*-th row of 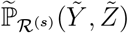 and the *j*-th row of 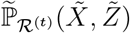, respectively. Specifically, for each cluster *i* ∈ *C*_*X*_, we extract the corresponding rows of 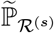 to obtain 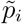; analogously, 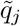 is constructed from rows in 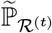 associated with cluster *j* ∈ *C*_*Y*_.

To identify matched clusters, we consider *n*_shuffles_ random permutations of the target cluster labels in *C*_*Y*_, indexed by *r* = 1, …, *n*_shuffles_. For each permutation *r*, clusters are matched in sequence using a greedy strategy that minimizes the *average Jensen-Shannon divergence* (AJSD):

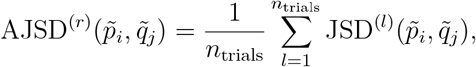

where in each trial *l*, equal-sized subsets are sampled from 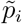 and 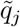 to mitigate potential imbalance between source and target clusters.

The **Jensen-Shannon divergence** between two discrete distributions *P* and *Q* is defined as:

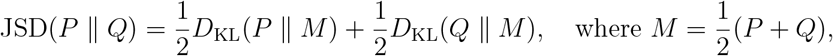

and *D*_KL_(· ∥ ·) denotes the Kullback-Leibler divergence.

For a fixed target cluster *j*, we select the best matching source cluster *k* by minimizing the AJSD:

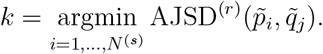

The match (*k, j*) is accepted if AJSD^(*r*)^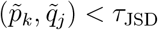, where *τ*_JSD_ is a predefined threshold (default: 0.45). If this condition is not satisfied, 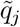 is considered unmatched—potentially corresponding to a novel cell type not observed in the source domain.

Within each permutation *r*, once a source cluster has been matched, it is excluded from further assignments in that permutation, enforcing an exclusive one-to-one matching constraint.

This matching process is repeated across *n*_shuffles_ permutations of *C*_*Y*_, producing a set of candidate matchings 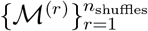. To determine the optimal matching configuration, we select the permutation *r*\* that minimizes the total divergence among all valid matches:

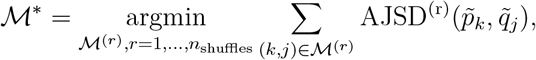

where the summation is restricted to matched pairs satisfying the AJSD threshold.

#### Rationale for Permutation-Based Matching

The permutation index *r* denotes an ordering of target clusters in *C*_*Y*_, e.g., [1, 2, 3, 4, 5] vs. [2, 3, 4, 5, 1]. Multiple permutations are evaluated for two key reasons: (i) *Exclusive Matching Enforcement* — once a source cluster is assigned, it cannot be reused in the same permutation, avoiding duplicate matches; (ii) *Optimization of Match Quality* — a fixed order may yield suboptimal results due to greedy assignments. For example, reordering clusters may better align with biological structure and minimize divergence. Therefore, we select the configuration that maximizes aggregate JSD, reflecting more separable and interpretable cluster correspondences.

In summary, the selected matching configuration ℳ* optimizes the divergence between matched clusters under exclusivity constraints, yielding high-confidence mappings that facilitate accurate and biologically meaningful label transfer.

### B.2 Algorithm

We provide the detailed algorithm of permutation-based matching across source and target clusters in Algorithm 2.

#### Complexity of Algorithm 2 (Permutation-Based Matching)

Let *N* ^(*t*)^, *N* ^(*s*)^ be #target and #source clusters, *T* = *n*_trials_, and *R* = *n*_shuffles_, with profile dimension *m*.

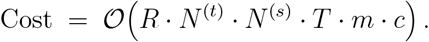

The parameter *c* represents the average number of rows utilized for Jensen-Shannon divergence (JSD) computations in each matching trial. This value is data-dependent, defined as 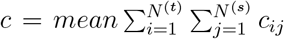, where *c*_*ij*_ = *min*(*nr*_*i*_, *nr*_*j*_), where *nr*_*i*_ and *nr*_*j*_ denote the row counts of i-th cluster of target data and j-th cluster of source data, respectively. The computational complexity for JSD calculation in a single trial is *O*(*mn*_*trials*_*c*). In worst-case scenarios, where all rows of either the source or target data might be processed, the theoretical time complexity of the matching algorithm scales as:

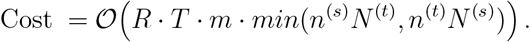

##### Algorithm 2 Permutation-Based Matching

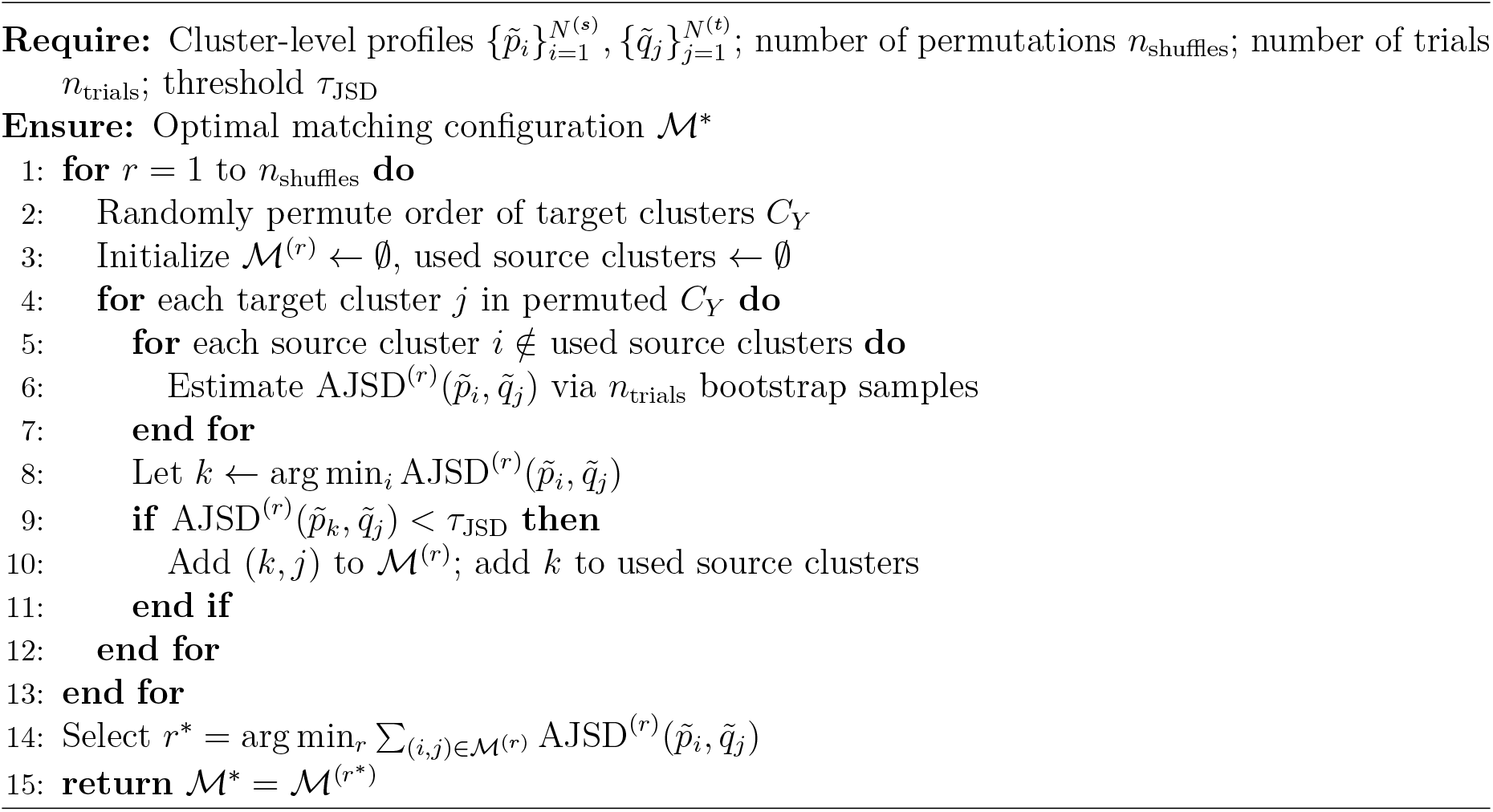

However, the practical complexity is reduced because previously matched clusters are excluded in subsequent iterations.

Notably, the matching algorithm’s time complexity is comparable to that of the CoCluster algorithm. Consequently, optimizing runtime efficiency necessitates balanced parameter configuration for both algorithms.

## C Parallel Computing Analysis for Algorithms 1 & 2

The co-clustering algorithm exhibits inherent parallelization potential in computing the function Q, which proceeds in two distinct stages. In the first stage, the adjusted distribution ℙ* is derived from the clustering results *C*_*Y*_ and *C*_*Z*_. Since ℙ* is uniquely determined by the cell index Y and feature index Z, this step is inherently free from data competition, enabling independent parallel computations across either the cell (Y) or feature (Z) dimensions. Subsequently, the second stage updates Q using ℙ and the precomputed ℙ*. With ℙ and ℙ* fixed at this stage, the value of Q depends solely on the current cell (or feature) and cluster cluster i. This independence eliminates data races and allows parallel execution across the dimensions of Y, Z (depends on currently upgrading *C*_*Y*_ or *C*_*Z*_), or cluster labels i. Both stages thus benefit from parallelization due to the absence of iterative dependencies and shared resource conflicts.

The matching function also exhibits inherent parallelization potential. First, since each permutation operation is entirely independent—with no permutation affecting the results of another—this process can be parallelized by distributing permutations across multiple threads for simultaneous computation. Furthermore, when calculating the Jensen-Shannon Divergence (JSD), each com-putation attempt operates independently of others, eliminating data competition or dependency constraints. This independence allows the JSD calculation phase to leverage parallel processing across threads. Both stages thus benefit from parallel execution due to their stateless, non-iterative nature. However, these two layers of parallelism are nested. While nested parallelism theoretically maximizes resource utilization, its practical implementation introduces significant scheduling overhead. Further more, some programming languages (e.g., `Matlab`) lack native support for hierarchical parallel task management. Consider that, the GuidedCoC framework restricts parallelism to a single level, JSD computation parallelization.

Current estimates indicate that 70–80% of operations are parallelizable. Under Amdahl’s law, this translates to a theoretical 2.3–2.9x speedup on a 4-core system.

## D Evaluation Metrics Details

Let *Q* be the predicted clustering result and *G* be the ground-truth labels. NMI measures the mutual dependence between *Q* and *G*. Let *I*(*Q*; *G*) be the mutual information, and *H*(*Q*) and *H*(*G*) be the entropies of *Q* and *G* respectively. NMI is computed as:

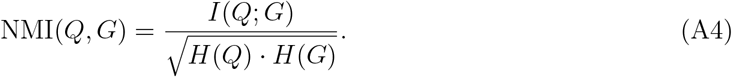

It ranges from 0 (no mutual information) to 1 (perfect correlation). Let *n*_*Q,i*_ be the number of samples in the *i*-th cluster of *Q, n*_*G,j*_ be the number of samples in the *j*-th class of *G*, and *n*_*ij*_ be the number of overlapping samples between these two groups. The ARI is computed as:

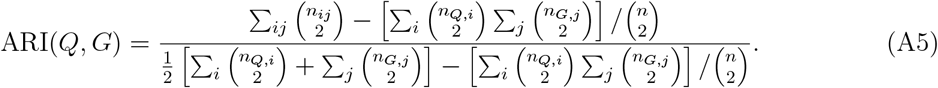

ARI corrects RI for chance, and ranges from -1 (random) to 1 (perfect match). Higher values of NMI and ARI indicates better clustering performance.

## E Dataset Details

We provide detailed descriptions of the four benchmark datasets used in our experiments:

- **Example 1 – PBMC Human Data:** The unpaired scRNA-seq data (3,709 cells) was collected from a healthy male donor (age 18–35) using the Chromium GEM-X Single Cell 3’ v4 platform (*10x Genomics*). This dataset is available at https://www.10xgenomics.com/datasets/5k_Human_Donor1_PBMC_3p_gem-x. The paired scRNA-seq + scATAC-seq multiome data (3,012 cells) was obtained from a healthy female donor using the Chromium Single Cell Multiome ATAC + Gene Expression platform. The datasets are available at https://www.10xgenomics.com/datasets/pbmc-from-a-healthy-donor-no-cell-sorting-3-k-1-standard-1-0-0.
- **Example 2 – E18 Mouse Brain:** The unpaired scRNA-seq data (11,843 cells) was derived from the cortex, hippocampus, and subventricular zone of an embryonic day 18 (E18) mouse brain using Chromium v3 chemistry. This dataset is publicly available at https://www.10xgenomics.com/datasets/10-k-brain-cells-from-an-e-18-mouse-v-3-chemistry-3-standard-3-0-0. The paired multiome data (4,481 cells) was generated from fresh E18 brain tissue using the ATAC + RNA multiome protocol. The datasets are available at https://www.10xgenomics.com/datasets/fresh-embryonic-e-18-mouse-brain-5-k-1-standard-1-0-0.
- **Example 3 – Mouse and Human Lymph Node:** The unpaired scRNA-seq data was collected from spleen and lymph nodes of an 8-month-old C57BL/6J mouse using the TotalSeq-C Mouse Universal Cocktail. We used the “Lymph node rep 2” sample. The dataset is publicly available at https://www.10xgenomics.com/datasets/Mixture-of-cells-from-mouse-lymph-nodes-and-spleen-stained-with-totalseqc-mouse-universal-cocktail. The paired dataset consists of human lymph node nuclei (14,645 high-quality nuclei) from a patient with diffuse small B-cell lymphoma, profiled using the Multiome ATAC + RNA platform. The datasets are available at https://www.10xgenomics.com/datasets/fresh-frozen-lymph-node-with-b-cell-lymphoma-14-k-sorted-nuclei-1-standard-1-0-0.
- **Example 4 – Pancreatic Islet:** The unpaired scRNA-seq data comes from human pancreatic tissues and is part of the GSE84133 dataset, which includes multiple donors and tissue types. The dataset is available at https://www.ncbi.nlm.nih.gov/geo/query/acc.cgi?acc=GSE84133. The paired multiome data was collected from healthy donors in the GSE200044 project, which profiled over 85,000 human islet cells from donors with various diabetic statuses. Only healthy donor cells were used in our analysis [35]. The dataset can be accessed at https://www.ncbi.nlm.nih.gov/geo/query/acc.cgi?acc=GSE200044.

## F Baseline Method Details

We briefly describe the baseline methods used for comparison:

- **MultiVI** [16] is a deep generative model based on variational autoencoders. It encodes scRNA-seq and scATAC-seq data into multivariate Gaussian latent spaces and aligns them by minimizing their divergence. Modality-specific decoders reconstruct the original observations, with training guided by a likelihood objective and adversarial regularization.
- **Cobolt** [14] models multimodal single-cell data using an LDA-inspired probabilistic framework. It assumes each cell is composed of latent biological categories with varying activation levels. A variational autoencoder is employed to infer a shared latent space and estimate the parameters governing the category distributions.
- **scMoMaT** [18] addresses mosaic integration by treating each data matrix as a relationship between cell and feature entities. It performs matrix tri-factorization to learn cell factors, feature factors, and an association matrix, optimizing reconstruction loss to capture shared and modality-specific signals.
- **Seurat v4** [8] integrates multimodal data using a weighted nearest neighbor (WNN) graph. It constructs KNN graphs for each modality, infers cross-modality similarities, and computes cell-specific modality weights to build a unified WNN graph, which is used for downstream tasks such as clustering.

### Implementation Details

To ensure a fair and representative comparison, we adopt the default hyperparameter settings provided by the original implementations for all baseline methods, which are commonly optimized for general-purpose performance [19]. All experiments are conducted on a Linux 64-bit system equipped with an Intel Core i7-14700K (water-cooled), 64 GB DDR5 5600 MHz RAM, an NVIDIA RTX 4090D 24 GB GPU, and a 1250W power supply.

## G More UMAP Visualizations

**Figure A1:**
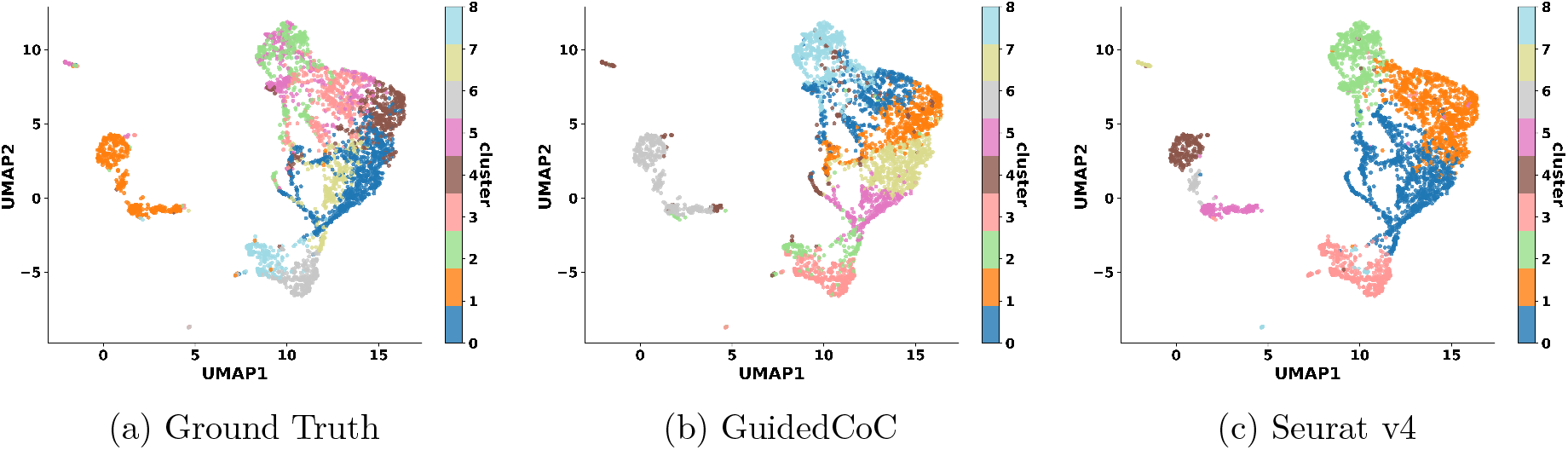
UMAP visualization of cells in the target scRNA-seq dataset ℛ^(*t*)^ (Example 2).

**Figure A2:**
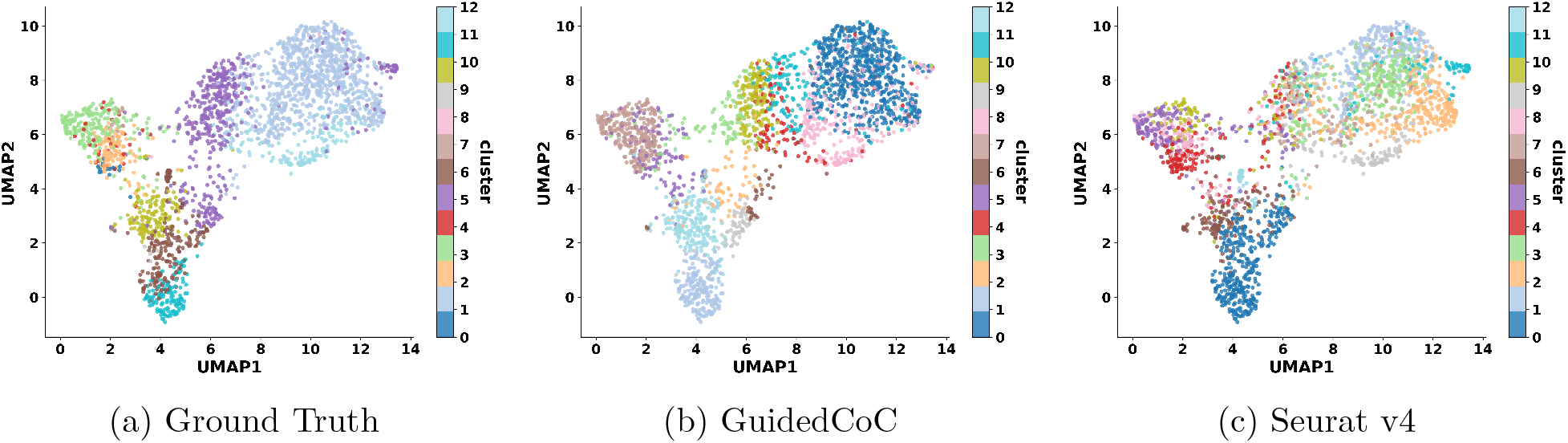
UMAP visualization of cells in the target scRNA-seq dataset ℛ^(*t*)^ (Example 3).

**Figure A3:**
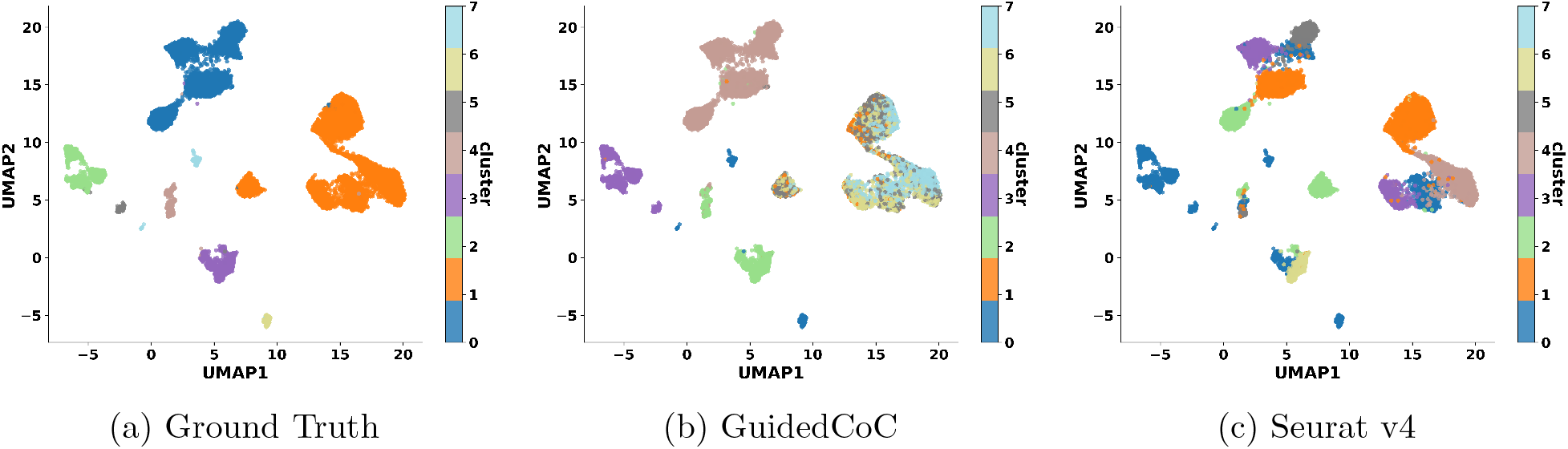
UMAP visualization of cells in the target scRNA-seq dataset ℛ^(*t*)^ (Example 4).

## H Details of Enrichment Analysis

**Table A1:** Selected enriched annotation terms for the gene lists in **Feature Clusters 1 and 2** of Example 1. Cluster 1 includes canonical T cell receptor genes such as *TRBV18, TRBC2, CD8A, CD8B*, and *CD247* ; Cluster 2 contains regulatory genes including *CD70, IL7, HLA-DPB1*, and *PDCD1LG2*.

| Category | Term | Count | % | Bonferroni P-value |
| --- | --- | --- | --- | --- |
| GOTERM_BP_DIRECT | Antigen processing and presentation | 11 | 5.64 | $4.51 \times 10^{-9}$ |
| UP_KW_CELLULAR_COMPONENT | T cell receptor | 14 | 3.44 | $2.76 \times 10^{-5}$ |
| GOTERM_BP_DIRECT | Positive regulation of T cell activation | 9 | 4.62 | $1.07 \times 10^{-6}$ |
| GOTERM_CC_DIRECT | T cell receptor complex | 15 | 3.69 | $5.68 \times 10^{-5}$ |
| GOTERM_BP_DIRECT | Adaptive immune response | 23 | 5.65 | $4.87 \times 10^{-2}$ |

**Table A2:**
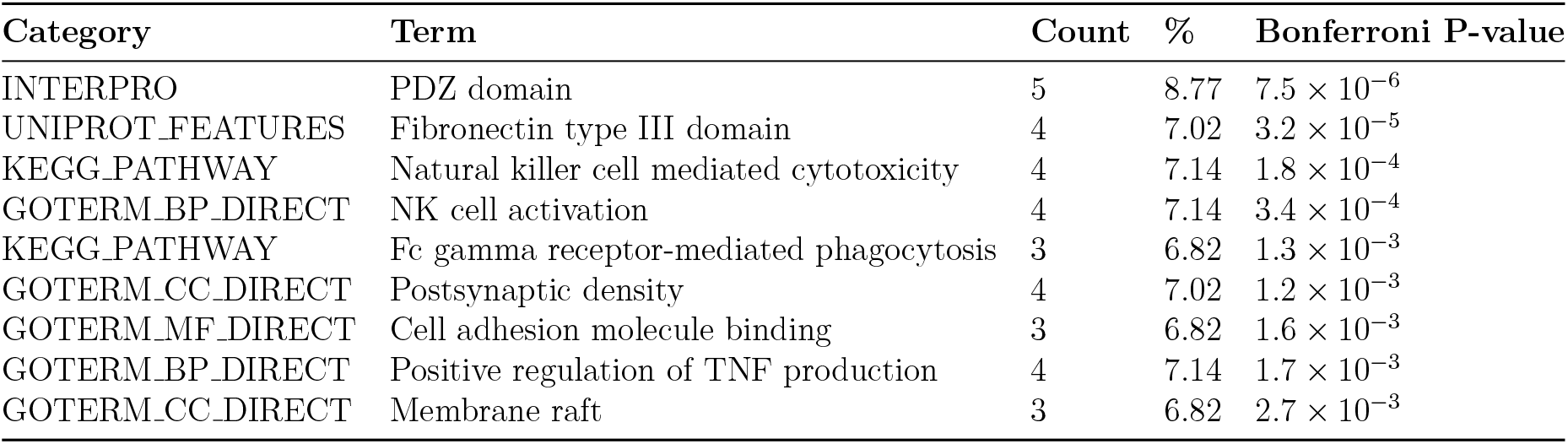
Selected enriched annotation terms for Feature Clusters 7–9. The gene lists include representative genes associated with immune functions such as *FCGR2B, DLG2, FCGR3A*, and the top pathways are shown.

| Category | Term | Count | % | Bonferroni P-value |
| --- | --- | --- | --- | --- |
| INTERPRO | PDZ domain | 5 | 8.77 | $7.5 \times 10^{-6}$ |
| UNIPROT_FEATURES | Fibronectin type III domain | 4 | 7.02 | $3.2 \times 10^{-5}$ |
| KEGG_PATHWAY | Natural killer cell mediated cytotoxicity | 4 | 7.14 | $1.8 \times 10^{-4}$ |
| GOTERM_BP_DIRECT | NK cell activation | 4 | 7.14 | $3.4 \times 10^{-4}$ |
| KEGG_PATHWAY | Fc gamma receptor-mediated phagocytosis | 3 | 6.82 | $1.3 \times 10^{-3}$ |
| GOTERM_CC_DIRECT | Postsynaptic density | 4 | 7.02 | $1.2 \times 10^{-3}$ |
| GOTERM_MF_DIRECT | Cell adhesion molecule binding | 3 | 6.82 | $1.6 \times 10^{-3}$ |
| GOTERM_BP_DIRECT | Positive regulation of TNF production | 4 | 7.14 | $1.7 \times 10^{-3}$ |
| GOTERM_CC_DIRECT | Membrane raft | 3 | 6.82 | $2.7 \times 10^{-3}$ |

